# Mapping the dimerization specificity of bZIP transcription factors in bread wheat

**DOI:** 10.1101/2025.02.12.637863

**Authors:** Raminder Kaur, Nishtha Sharma, Prateek Jain, Mamta Verma, Ashish Apkari, Vikas Rishi

## Abstract

Dimerization is pivotal in target-binding interactions among transcription factors (TFs), including bZIPs, bHLHs, and many zinc finger proteins. The bZIP family comprises an α-helix class of TFs that features a basic region binding to the major groove of double-stranded DNA (dsDNA), followed by a leucine zipper motif that facilitates dimerization through coiled-coil structures, forming homo- or heterodimers. In allohexaploid bread wheat (*Triticum aestivum* cv.), bZIPs and their homeologs utilize dimerization motifs to create specific bZIP pairs with distinct regulatory functions. The dimerization pattern of bZIPs is essential for gene regulation. Following a genome-wide analysis, we identified 265 bZIP TFs in bread wheat. By applying criteria from previous studies to evaluate bZIP dimerization across humans, Drosophila, and Arabidopsis, we categorized bZIP TFs into different groups based on their predicted dimerization propensity. Amino acid sequence analysis of wheat bZIP TFs led to several conclusions: unlike Drosophila and humans, wheat bZIPs are longer, with some reaching lengths of up to 14 heptads. Based on the amino acids at the a, d, g, and e positions in each heptad, wheat bZIPs are predominantly predicted to form homodimers. However, the mechanisms underlying wheat bZIP dimerization specificity and stability, particularly in plants, remain poorly understood. To experimentally validate the predicted dimer assembly, eight key bZIP TFs containing the bZIP domain were cloned and expressed for structural analysis. Using purified bZIP proteins, thermal denaturation studies demonstrated a change in dimerization stability as the length of the leucine zipper varies. In summary, this study enhances our understanding of the role of leucine zippers in determining the specificity and stability of bZIP dimers, which subsequently influence DNA binding and gene regulation in plants.

## 1. Introduction

### 1.1. bZIP Transcription factors (TFs)

bZIP TFs serve as the primary mediator of environmental signals that ultimately result in gene expression. DNA-binding class of TFs interact with transcription factor complex to recruit or block the access of RNA polymerase to the promoter region of the gene, and hence trigger the gene regulation process (Krylov *et al*. 1998). Several regulatory DNA-binding transcription factors are classified based on the DNA-binding domains recognized by them such as bZIPs, bHLHs, MYBs, NACs, etc (Riechmann and Ratcliffe 2000). Many DNA binding proteins employ dimerization as an essential feature for target-binding interactions (Baxevanis and Vinson 1993).

The term bZIP comes from the basic amino acid-rich N-terminal region and the C-terminal region containing amphipathic amino acids (Krylov *et al*. 1998). Although a few bZIPs have been well characterized, the majority of the bZIP family members remain to be further explored. bZIPs are crucial for various aspects of plant development and defense (Jakoby *et al*. 2002). The bZIP proteins contain a continuous α-helix, including the basic region, which is essential for binding to the major groove of double-stranded DNA. This is followed by the C-terminal leucine zipper motif, which facilitates dimerization through a coiled-coil structure (Acharya *et al*. 2002). In bZIPs, the leucine zipper motif plays a crucial role in the functional activity of various transcriptional regulatory proteins, particularly in dimerization (Moitra *et al*. 1997). bZIPs bind to specific sequences of DNA. Plant bZIP transcription factors like to bind to the common ACGT component sequences A-box (TACGTA), C-box (GACGTC), and G-box (CACGTG) (Foster *et al*. 1994). In addition, they have been reported to bind to non-core ACGT DNA motifs (Foster *et al*. 1994; Kusano *et al*. 1995).

Researchers initially divided the members of the bZIP TF family in *Arabidopsis* into ten groups based on common domains (Jakoby *et al*. 2002). Further, the bZIPs were classified into 13 groups (A-M) (Dröge-Laser *et al*. 2018). Vinson and co-workers divided the Arabidopsis bZIPs into different groups based on their predicted dimerization properties (Deppmann *et al*. 2004). However, a comprehensive classification of plant bZIPs based on dimerization specificity remains lacking, especially in complex plant genomes like bread wheat (Triticum aestivum).

### 1.2 Structural signatures determining the dimerization specificity in bZIPs

Dimerization has emerged as an integrated property manifested by DNA binding proteins (Vinson *et al*. 1989; Pathak and Sigler 1992). Dimerization involves the potential for the monomers to form dimers with the associated mechanisms, which enhances their function for the incremental biological control (Amoutzias et al., 2006, 2008). Newman and Keating have shown that, in Humans and Yeast, about 15% of all possible interactions occur among bZIP protein (Newman and Keating 2003).

The bZIP structure comprises 60–80 amino acids, including an N-terminal basic DNA binding region and a C-terminal adjacent leucine zipper region (Ellenberger *et al*. 1992; Deppmann *et al*. 2004). The DNA binding region is abundant with basic amino acids and has a conscious N-X_7_-R/K motif in its DNA-binding sequence (Landschulz *et al*. 1988). The bZIP monomer comprises a structural repeat of amino acids (Krylov *et al*. 1994). Each structural repeat is termed a heptad consisting of seven residues designated with alphabets (*a,b*,*c*,*d,e*,*f*, and *g)*(Acharya *et al*. 2002; Vinson *et al*. 2006). The ‘a’, ‘d’, ‘e’ and ‘g’ are prime positions in the leucine zipper interface and are responsible for the dimerization stability and specificity (Acharya *et al*. 2002, 2006; Deppmann *et al*. 2004; Vinson *et al*. 2006).

The amino acid residues at ‘*a*’ and ‘*d*’ positions form the hydrophobic interface on the similar edge of the α-helix (Pathak and Sigler 1992). These residues interact inter-helically and stabilize the structural dimer (Vinson *et al*. 2002, 2006). Leucine is frequently present in the *’d*’ position and stabilizes the dimer more effectively than other amino acids (Moitra *et al*. 1997). In this configuration, the hydrophobic core formed between the non-polar surfaces of two leucine zippers causes them to dimerize (Vinson *et al*. 2002, 2006; Deppmann *et al*. 2006).

Electrostatic interactions are generated by the charged amino acids at the ’g’ and ’e’ positions in a heptad (Krylov et al. 1994). The ‘g’ and ‘e’ interactions are indicated as g ↔ e ′ where the prime (′) indicates a residue on the second interacting α-helix of the dimeric leucine zipper. g-e ′ interactions between oppositely charged residues encourage homodimerization (Deppmann et al. 2004). The g-e ′ interactions between likely charged residues are repulsive and encourage heterodimerization. In plant bZIPs, similar positioning of crucial amino acids is observed as in humans. Asparagine occurs at ‘*a*’ position in the second heptad of all human bZIPs, which results in homodimerization typically (Jakoby *et al*. 2002). In humans, 32% of Asparagine occurs in the second heptad at the ‘*a*’ position, whereas it is about 68% in *T. aestivum* in the present study.

The leucine zipper region comprises hydrophobic amino acids such as leucine, isoleucine, and valine (Vinson *et al*. 1989). It has also been noted that distinct heptads vary substantially in length, and the influence these length differences have, on the functional features, is a topic of active research. Understanding these structural determinants is critical for predicting bZIP dimerization behavior, particularly in complex plant genomes like wheat, where expanded bZIP families may contribute to diverse regulatory pathways. Heterodimerization is common in mammalian systems, where bZIPs like the Jun family interact with FOS members to form oncogenic complexes (Vinson *et al*. 2002). No similar system is reported in plants. To address this gap, we looked at the ability of the bread wheat bZIP sequences to form homo-or heterodimers, which may help to uncover DNA sequences that these bZIPs recognize and, by logical extension, novel genes that they may regulate.

## 2. Materials and methods

### 2.1. Plant materials

Bread wheat (Triticum aestivum) variety Chinese Spring (CS) cultivar (cv.) was grown in the experimental fields of the BRIC-National Agri-Food and Biomanufacturing Institute, Punjab, India. First day after anthesis (DAA) was considered when anthers appeared from the central portion of the spike. After the anthesis appeared, the plants were tagged for sampling at different stages of seed development such as 0,7,14,21 and 28 days after anthesis (DAA). Seed samples from spikes at different developmental stages i.e. at 7, 14, 21 and 28 DAA which corresponds to stage Z-72, Z-75, Z-78, Z-82, respectively, according to Zadoke scale were collected and stored at -80°C (ZADOKS *et al*. 1974). Tissue samples such as leaf, flag leaf, stem and root were also collected, frozen in liquid nitrogen (N_2_) and stored at -80°C until further use.

### 2.2. Cloning of bread wheat bZIPs

For the zipper region of candidate bZIP genes, the primers were designed for the cloning experiments. Different parameters were considered including Tm, GC content, and hairpin loop structure using Oligo Calc tool. Zipper-region of bZIPs were amplified as BamHI-HindIII fragment using Taq DNA polymerase. Quality of PCR products was checked on 1.2% agarose gel. For ligation of amplified bZIPs into pT5 protein expression vector, conventional cloning was used. All cloned fragments have T7 tag in their N–terminus region. Positive clones from pT5 vector were confirmed by Sanger-dideoxy sequencing and were attempted for sub-cloning in expression vector pET23b+. The expression vector was considered because of higher expression of bacterial recombinant proteins. Details are described in supplementary material S1 file for the methodology of recombinant protein expression, downstream processing, purification and quantification.

### 2.3. Multiple alignment for prediction of dimerization specificity

Eight different TabZIPs (Embp1, TabZIP1, TaABI5 well reported in earlier studies and TraesCS4B02G320400, TraesCS5A02G237200, TraesCS7A02G398400, TraesCS1A02G072600 and TraesCS7D02G518100 as novel bZIPs) from hexaploid wheat were selected on the basis of their expression which have potential role in seed maturation and abiotic stress. Amino acids sequences of seed-specific bZIPs were aligned using the multalin sequence alignment tool. The presence of an invariant asparagine (**N**) residue in basic region of bZIPs was used for alignment of protein sequences. The N-terminal boundary for bZIP is conserved due to presence of basic amino acids while to establish C-terminal boundary, following four criteria were used: **(i)** Presence of Proline (P) or **(ii)** two constitutive Glycines (G), that disrupts α–helical structure **(ⅲ)** repetition of leucine at ‘d’ position in leucine zipper and **(Ⅳ)** Presence of charged amino acids at ‘e’ and ‘g’ position (Deppmann et al., 2004). After alignment, the N–terminal and C–terminal boundaries were defined and amino acids responsible for dimerization were color coded according to previously published work (Fassler et al., 2002; Vinson et al., 2002; Deppmann et al., 2004).

### 2.4. Protein secondary structure analysis using Circular Dichroism

For CD, all eight protein samples used, were of highest purity. The presence of any insoluble aggregates can distort the shape and magnitude of CD spectrum that will decrease the signal/noise ratio (Kelly *et al*. 2005). For wavelength and thermal denaturation studies experiments were performed in standard CD buffer (12.5mM KPO_4_ buffer pH 7.4, 150 mM KCl and 1M DTT). All circular dichroic studies were carried out using Biologic CD spectrometer (Biologic, France). Before CD studies, samples were heated to 65°C for 15 min and allowed to cool at room temperature. For structural studies, 2 mΜ (dimer) proteins were used. Wavelength scans of proteins were carried at 6°C and samples were scanned from 200-260 nm. Average of 5 scans for each protein sample was taken. For thermal denaturation studies protein samples were heated from 6-85°C at the rate of 1°C/min while change in ellipticity was measured at 222 nm. All TabZIP proteins melted with the well-defined sigmoidal transition curves. All such curves were fitted according to (equation 1) assuming denaturation to be a two-state type.

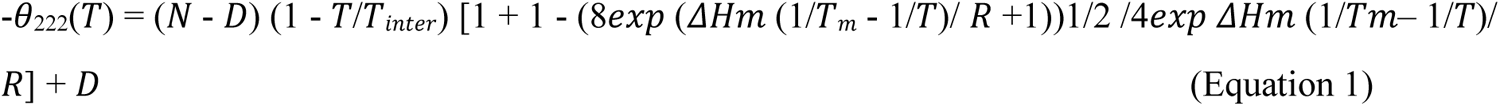

where -θ222 (T) were CD signals in millidegree (mdeg) at 222 nm at any temperature T in °C, N is the ellipticity of the α-helical coiled coil dimer at 0°C extrapolated from the linear-dependencies of CD signals at low temperature range defined as pre-transition region, D is the ellipticity at high temperature extrapolated to 0°C and referred to as post-transition baseline where dimeric protein molecules melt to unhelical monomers. Transition region between pre-and post-transition region has monomer and dimer bZIPs in equilibrium. T inter is the temperature at which the linear temperature-dependencies of dimer and monomer molecules intersect, and Ris the gas constant. Fitting of thermal denaturation curves gave values of midpoint of thermal denaturation (Tm), and enthalpy change at Tm (ΔHm). Constant-pressure heat capacity change (ΔCp) was calculated using Tm versus ΔHm plot for all the possible bZIP dimers. A linear fitting of these data points with negative slope gave the value of ΔCp. This along with Tm, and ΔHm values were used to calculate ΔGDi, Gibbs free energy of dimerization at 25°C for each dimer by using the following form of Gibbs Helmholtz equation (Equation 2).

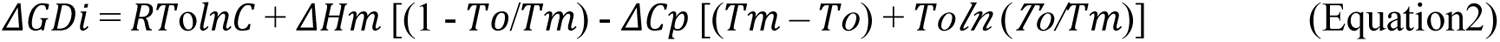

where *To* is any temperature in °C and C is total protein concentration.

### 2.5. Identification of amino acid variations and leucine zipper length

Being a polyploid plant, bread wheat TabZIPs were aligned (Supplementary figure 1). While aligning the sequences of the TabZIPs, some were identified as exceptionally longer and with one or more amino acid variations within the homeolog copies. TraesCS4B02G320400, TraesCS5A02G237200 out of eight TabZips, were chosen for the biochemical characterization devoted by their structural features. Presence of specific amino acids, owing to the distinct structural properties is leading to the stable helical structure formation in the bZIPs. Identification further led the basis to analyse their α-helical structure characterization and thermal stability behavior using CD spectroscopy.

### 2.6. MALDI-TOF-MS protein profiling

10 mg of sinapinic acid (SA) stock was mixed with the sample in equimolar ratio and was dried on a microtiter plate format 384 well target made out of full metal (MTP) plate. The dried layer was analyzed by a ABSCIEX TOF/TOF 5800 MALDI instrument in the mid-mass linear positive mode.

One microliter of the sample/matrix solution was spotted onto the MALDI target plate MTP 384 and left to crystallize at room temperature. All protein samples were analyzed in a random order to minimize variability and systematic errors. ABSCIEX TOF/TOF 5800 MALDI-TOF mass spectrometer (SCIEX, USA) was used to perform MS analyses in the mid mass linear positive mode. Positively charged ions were detected in the m/z range of 9000–27,000 Da and 1000 shots were accumulated per one spectrum. The MS spectra were externally calibrated using the mixture of following Sigma protein calibration standards such as ProteoMass^™^ Bradykinin Fragment 1-7, ACTH Fragment 18-39, Insulin chain B oxidized, Angiotensin II, Insulin, P_14_R etc. at the ratio of 1:5.

The average mass deviation was less than 100 ppm. The matrix suppression mass cut off was m/z 700 Da. The following ion source parameters were used: ion source 1, 25.09 kV; ion source 2, 23.80 kV. Other settings for MALDI-TOF MS analysis were as follows: pulsed ion extraction, 260 ns and lens, 6.40 kV. In-built software TOF-TOF series explorer software (Ab Sciex, U.S.A) was applied for the acquisition and processing of the spectra. Each sample was analyzed in three repetitions. Inter-day and intra-day reproducibility of the applied procedure was evaluated in our study.

## 3. Results

### 3.1. Chromosomal localization of bZIP transcription factors of bread wheat

All the TabZIPs were attempted for the localization on the chromosomes covering around 91% of the whole genome as shown in Figure 1. *bZIP* genes distribution on chromosomes varies greatly and is uneven. For example, chromosomes 5 and 7 have the most TabZIP (24% and 13% respectively) but these TFs are less on chromosome 6 with 10 % distribution (Figure 1 (b)). Chromosome 2 and 3 contained twenty-six members (12% and 14%). Further, chromosomes 1 and 4 have 11 TabZIPs, with a 11% share in the genome. Interestingly, most of the genes are present near the ends of the chromosomes suggesting a dynamic gene expression trend opted by the *TabZIP* genes.

**Figure 1.**
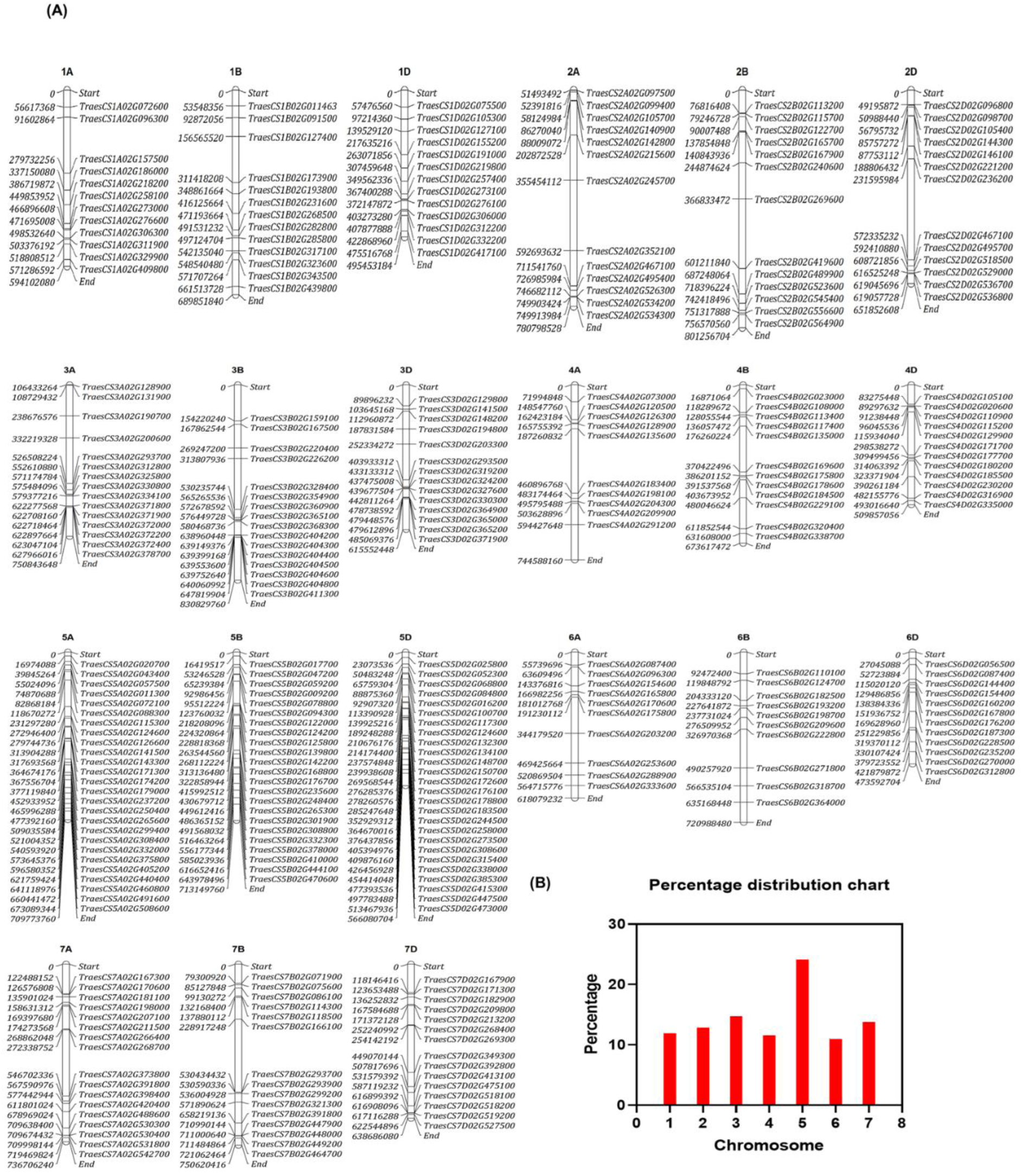
Genome wide distribution of TabZIPs on wheat chromosomes. **(A)** Distribution of bZIP proteins is represented on wheat genome on the basis of their chromosomal locations. Each chromosome is represented with black color (B) panel shows the percentage distribution of bZIPs on all chromosomes.

### 3.2 Genome-wide annotation of dimerization properties of bZIP transcription factors in bread wheat

A structure-based search of the wheat transcription factors databases such as ***expVIP*** and ***EnsemblPlants*** has identified 265 bZIP TFs in hexaploid wheat. To predict bZIP dimerization specificity in wheat, structural rules established for the human (Vinson *et al*. 2002), drosophila (Fassler *et al*. 2002), and *Arabidopsis* (Deppmann *et al*. 2004) were considered and are as follow-

i. length of the leucine zipper,
ii. placement of asparagine or a charged amino acid in the hydrophobic interface, and
iii. presence of interhelical electrostatic interactions.

Though literature is scarce but predicted dimer partner choice is not random in these bZIPs, instead dimerization specificity is defined by ‘g’ and ‘e’ positions followed by the presence/absence of asparagine at the ‘a’ position in a heptad. Following the above-entrenched criteria, we have divided the bread wheat bZIPs into eleven groups. Supplementary figure 1 shows the amino acid alignment of 265 wheat bZIPs. The specifics presented below is in accordance with the observations. Figure 2 shows the color-coded interactions between the candidate TabZIPs used in the study. We have colored-coded the interactions based on the type of amino acids in the ‘a’, ‘d’, ‘e’, and ‘g’ positions in a heptad. Four colors have been used for g↔e′ interacting pairs. Green is for the attractive basic-acidic pairs (R (arginine) ↔ E (Glutamic acid) and K (Lysine) ↔ E), orange is for the attractive acidic-basic pairs (E↔R, E↔K, D (Aspartic acid) ↔R, and D↔K), red is for repulsive acidic pairs (E↔E and E↔D). Blue is for repulsive basic pairs (K↔K and R↔K), etc. We have colored the residue red for acidic and blue for basic if just one of the two amino acids in the g↔e ′ pair is charged. It is black if the ‘a’ or ‘d’ position is polar and purple if either is charged. The following features of the specific amino acid residues in the heptads, have been postulated below:

**Figure 2.**
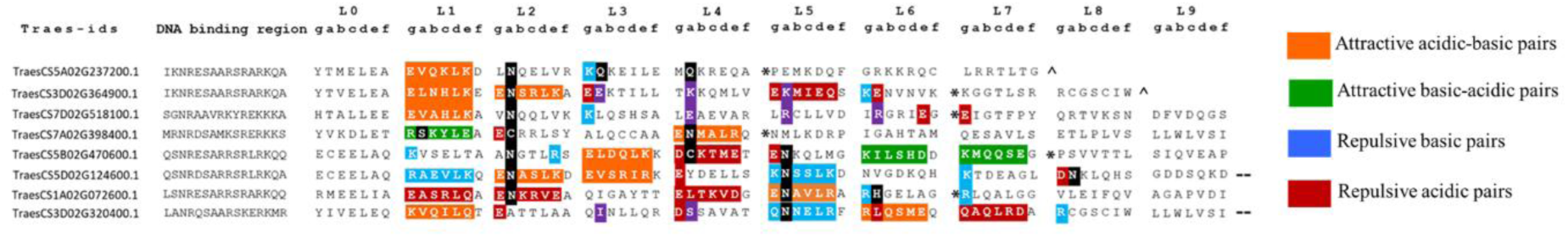
Amino acid sequence of all identified TabZIPs in bread wheat. Protein sequences are aligned based on the similarity in sequence. Second column contains Traes ids. Third column has n-terminal region of bZIP domain and c-terminal leucine zipper region is divided into heptads (a,b,c,d,e,f and g). Primarily, the placement of N at ‘a’ position was considered for classification and it was coloured black at the respective ‘a’ position. Secondarily, for the g-e’ interactions, we used four colours (provided on the below right side). The attractive acidic-basic pairs (E↔R’, E↔K’, D↔R’, D ↔K’) were coloured orange, and the attractive basic-acidic pairs (R↔E, K↔E’) were coloured green. We coloured the repulsive basic pairs (K↔K’, R↔K’) with blue and repulsive acidic pairs (E↔E’, D↔D’, E↔D’) with red. We have coloured the polar residues black at ‘a’ and ‘d’ position and charged residues at ‘a’ and ‘d’ positions were coloured purple. The candidate genes have been divided into five groups. We have used (**\***)symbol for potential helix break, due to the presence of proline or glycine. Symbol (**^**) denotes the natural end of a bZIP protein, and (--) shows that it is a longer protein and we have only shown the 9 heptads in the coloured figure.

#### 3.2.1 Eleven bZIP sub-groups in *T. aestivum*

Like other polyploid crops, sugarcane and cotton, Triticum aestivum, is a hexaploid crop. It contains 42 chromosomes with contributions from three, i.e., A, B, and D genomes and six allele copies in total. It gives three homoelog triad copies of every bZIP in hexaploidy wheat T aestivum. Here, group 1, the largest sub-family, is classified based on the presence of an N at the ‘a’ position in the 2^nd^ heptad, comprising 86 members. Attractive acidic-basic and attractive basic-acidic g↔e′ interactions in the 1^st^, 2^nd^, 5th and 6^th^ heptads. Additionally, repulsive g↔e′ interactions are present in 1^st^, 3^rd^, and 5^th^ heptads favoring both the aspects of homo- and hetero-dimerization between the same TabZIP protein or within the sub-family. Furthermore, single repulsive acidic and basic amino acids are present at the ‘g’ and ‘e’ positions, individually deciphering the heterodimerization outcomes from this family member protein. The 1^st^ heptad contains attractive g↔e′ acidic-basic interactions in this group. The presence of positively and negatively charged residues at the ‘g’ and ‘e’ positions instigates the electrostatic interactions. Its zipper (dimerization) region ends after the 6^th^ heptad because of the presence of P, a helix breaker.

Group 2 is classified based on the presence of N at a position of 4^th^ heptad. It also has glutamic acid at ‘e’ position in 2^nd^ heptad of this group member. It is composed of a single bZIP protein which is nine heptads long.

Group 3 consists of 15 bZIPs. It is characterized by N at ‘a’ position in the 2^nd^ and 4^th^ heptad and attractive basic-acidic pairs (R↔E and K↔E) in heptads 1 and 2. Single repulsive acidic and basic amino acid residues in 1^st^, 2^nd^, and 3^rd^ heptads are also present. This group contains heptad lengths ranging from 11 to 14. Concurrently, S is present at the ‘*a*’ position in some of the members of this group in the 1^st^ heptad. The unique placement of N in the ‘*a’* position of the 2^nd^ and 4^th^ heptads will favor homo-dimerization.

In Group 4, there are 16 members. The attractive *g*↔*e*′ pair in the 3^rd^ heptad (except one) should favour homodimerization. There are repulsive interactions in 7^th^ heptads (except three members) that may promote heterodimerization. The presence of N in ‘*a’* position of the 5^th^ heptad should favour homodimerization within the group members. Incomplete and/or repulsive basic pairs in the 1^st^ and 3^rd^ heptads indicate likely heterodimerization with acid zipper proteins. The presence of attractive acidic-basic *g*↔*e*′ interaction in 5^th^, 6^th^, and 9^th^ heptads and attractive basic-acidic interaction in 7^th^ heptad should lead to homodimerization., but will not impact much because of repulsive *g*↔*e*′ acidic and basic amino acid abundance in initial heptads. These initial heptads are decisive for dimerization.

Group 5 comprises of 21 bZIP members. Placement of N in ‘*a’* position of 2^nd^ and 5^th^ heptad and an attractive *g*↔*e*′ interaction in the 2^nd^ heptad indicates homodimerization. The presence of repulsive *g*↔*e*′ acidic and basic amino acid abundance in 2, 3, 4, 5 and 6 heptads, makes it the most complex group in defining dimerization fate.

Group 6 with 6 bZIP members having N at ‘a’ position in 2^nd^ and 4^th^ heptads. It has attractive basic-acidic *g*↔*e*′ pairs in 1 and 2^nd^ heptads and attractive basic-acidic *g*↔*e*′ pairs in 4^th^ heptads, suggesting homodimerization.

Group 7 has 20 bZIP members. The placement of N in the ‘*a’* position of the 5^th^ heptad indicates homodimerization, but there is a likelihood of heterodimerization because of the presence of repulsive acidic and basic *g*↔*e*′ pairs in 2, 3, and 7^th^ heptads.

Group 8 has 56 bZIP members. It is expected to homodimerize due to the presence of N in the ‘*a’* position of the 1^st^ and 5^th^ heptads and two attractive *g*↔*e*′ pairs in the 2^nd^, 3^rd^,5^th^, 6^th^, and 7^th^ heptads. Similarly, the presence of repulsive acidic and basic *g*↔*e*′ interactions in almost1,2,3,4,5,6 and 7 heptads of around 17 group members may lead to heterodimer formation with other groups.

Group 9 has 33 bZIP members. Attractive *g*↔*e*′ interactions in the 5^th^, 6^th,^ and 9^th^ heptads and the presence of N in the ‘*a’* position of the 5^th^ and 8^th^ heptad suggests homodimerization. The presence of charged amino acids in ‘*g’,* and *‘e’* position in the 1^st^, 2^nd^, 4^th,^ and 7^th^ heptads followed by attractive *g*↔*e*′ pairs in the 4th and 5th heptads may lead to the heterodimerization with other groups.

Like group 2, the group 10 is the smallest groups with one member strength with an unpredictable dimerization behaviour.

Finally group 11 has six bZIP members. It has a mix of attractive and repulsive *g*↔*e*′ pairs but has N at ‘*a’* position in the 2^nd^, 5^th^ and 9^th^ heptads, encouraging homodimerization.

### 3.3 Features of a, d, g, and e positions amino acids in the leucine zipper domain of wheat bZIPs

The interaction interface reveals four critical sites in bZIP heptads, which play a pivotal role during the dimerization event. We have compared the features of these positions in *T. aestivum* with the data available from other reference species such as *Z mays*, *O sativa*, *H vulgare*, and *A thaliana* in Figure 3 (Deppmann *et al*. 2004; Nijhawan *et al*. 2008; Wei *et al*. 2012; Zhong *et al*. 2021) . Amino acids in the ‘a’ and ‘d’ positions of the leucine zipper are typically hydrophobic, with various amino acids in the ‘a’ position and predominately leucine in the ‘d’ position. An exception is a 2^nd^ heptad ‘a’ position, which contains N in most homodimerizing bZIP proteins.

**Figure 3.**
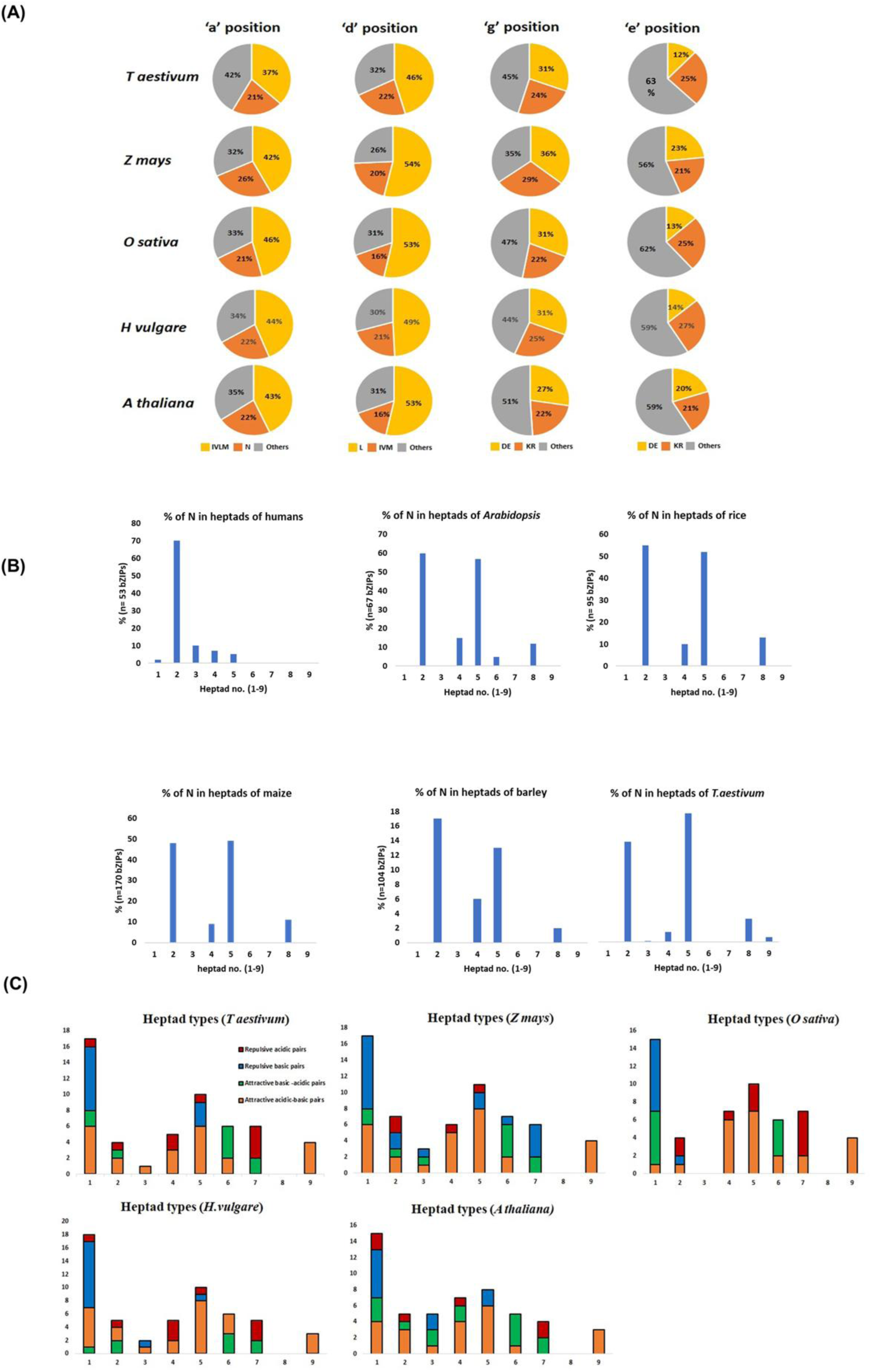
Features of ‘a’, ‘d’. ‘e’ and ‘g’ positions in heptads of TabZIPs. **(A)** Pie chart representing the frequency of occurrence of amino acids in all the *a,d,e and g* positions of heptads in the leucine zipper in *T. aestivum* bZIP family (B) Histograms presenting the dimerization details deduced solely from the presence of N residue at ‘a’ position in the leucine zipper region of *T. aestivum* and other reference species such as *H. sapiens, A. thaliana, O. sativa, Z. mays* and *H. vulgare*. **(C)** Histogram showing the frequency of attractive or repulsive g↔eʹ pairs per heptad for bZIP proteins in bZIP domains of T aestivum, Z mays, O sativa, H vulgare and A thaliana

It is found that 21% of *T aestivum* bZIPs contain N at ‘a’ position suggesting homodimerization as N prefers to interact with N in the second monomer participating in the homodimerization. In addition, *T aestivum* has the highest concentration of hydrophobic amino acids among all plant species, with 37% of the ’a’ position occupied by I, V, L, and M. It may prevent the higher order oligomerization, emphasizing the importance of ‘a’ positions equivalent to ‘g’ and ‘e’ positions in deciding the interaction fate. On the other hand, N at the ‘a’ position is highest in *Z mays* and suggests its inclination in forming homodimers. L at the ‘d’ position stabilizes the coiled-coil of bZIPs more effectively than any other amino acid in the hydrophobic interface. In a few cases, it is replaced by other hydrophobic aliphatic and aromatic amino acids, such as I, V, M, and F. In *T aestivum*, 46% of ‘d’ position has leucine, which is less than *A thaliana* (53%), *O sativa* (53%), *H vulgare* (49%), and *Z mays* (54%), suggesting that the gene families have undergone evolutionary modifications throughout the hybridization process. The 22% occurrence of other amino acids at the ‘d’ position, such as I, V, and M, is highest in *T aestivum*.

All TabZIPs contain charged and long side-chain amino acids such as D, E, R, K, and Q in heptads. The presence of charged amino acids at the ‘e’ and ‘g’ positions is crucial. They augment (+ ↔ - or - ↔ + electrostatic interactions) or diminish (- ↔ - or + ↔ + interactions) the dimer formation among bZIPs depending on the types of amino acids present at these positions. In wheat, acidic amino acids D/E, with 31 %, are more common in the ’g’ position than in Arabidopsis (27%) and less common than in *Z mays* (36%). On the contrary, the basic amino acids such as K/R are present in 24% of cases, the highest among all plant species except Z *mays* (29%). Besides, an abundance of D/E residues at the ‘e’ position is scant in *T aestivum* (12%), and is less than *Z mays* (23%), *O sativa* (13%), *A thaliana* (20%), and *H vulgare* (14%). Furthermore, *T aestivum* has a 25% distribution of K/R residues, higher than *O sativa* (22%) and *A thaliana* (22%).

### 3.4 Prediction of dimerization properties of candidate TabZIPs in wheat compared to the reference plant species

We have predicted the dimerization properties in the reference plant species (Oryza sativa, Zea mays, Hordeum vulgare, and Arabidopsis thaliana) used for the study with the entrenched structural rules of dimerization discussed in the previous section 3.1.1. Candidate TabZIPs and some putative bZIPs with a strong expression in different seed development stages, we have prepared the Supplementary figure 2 for *A thaliana, O sativa, Z mays,* and *H vulgare.* Furthermore, the dimerization properties of each species were compared with *T aestivum* data (Supplementary figure 2 (A)) to understand the differences between the plants used in the study and the significance of using *T aestivum* as a candidate plant species.

The dimerization specificity in the bZIPs is governed by a conserved structural rules (Vinson *et al*. 2002). Therefore, we have conceived the idea of classifying the candidate TabZIPs based on the amino acid present in each heptad. The amino acid sequence of these bZIPs in bread wheat is shown in the color-coded Supplementary figure 2 (A), aligned to show the dimerization potential. The N-terminal boundary is similar for all bZIP proteins defined by the presence of the clusters of basic amino acids. Further, conserved N residue in the basic region of all the bZIPs was used to align the bZIP sequences. We employed the same structural criteria used earlier for the dimerization prediction of the entire group in Supplementary figure 1.

The wheat bZIPs’ leucine zipper region contains electrostatic interactions that are both attractive and repulsive. These interactions are shown in colored boxes and are used to predict the dimerization potential of a specific bZIP protein. Observations are consistent with the bZIP families identified in Humans, Drosophila, Arabidopsis, and cereal crops like rice and maize (Fassler *et al*. 2002; Jakoby *et al*. 2002; Vinson *et al*. 2002; Nijhawan *et al*. 2008; Wei *et al*. 2012). The distinct combination of amino acids at the a, d, e, and g positions creates a new set of structural guidelines for determining the species-specificity of leucine zipper dimerization (Deppmann *et al*. 2004) bZIP proteins regulate the function of target genes by forming homo- or heterodimers. To determine the molecular basis of heterodimer formation between TabZIP proteins, we analyzed their propensities to form dimeric complexes.

#### 3.4.1 Homo- and Heterodimerization potential within the candidate TabZIPs

The key step for each bZIP in every species is determining the likelihood of selecting the suitable dimerization partners, which ensures the success of the dimerization process. Figure 2 illustrates the homo- and heterodimerization properties of the candidate TabZIPs analyzed in this study, highlighting the possible interaction patterns among them.

TraesCS5B02G470600 (EmBP1) TF has three attractive acidic-basic pairs and four repulsive acidic or basic pairs. Out of the repulsive pairs, three incomplete and one complete acidic pair are present. Again, in the second heptad of next pair TraesCS5B02G470600 (EmBP1) with TraesCS7D02G518100, TraesCS1A02G072600, and TraesCS3D02G364900 (ABI5), there is anticipatory formation of heterodimers between repulsive basic amino acid residues K↔K’, K↔Q’, and K↔K’ at g↔e’ positions. Following the dimerization trend between the TraesCS5B02G470600 and TraesCS7D02G518100 having K↔K’ at g↔e’ positions in the fourth heptad of their sequences.

Secondly, TraesCS7A02G398400 being the weakest TabZIP possesses incomplete dimerization potential due to lack of charged amino acid residues at the ‘g’ and e positions. There is little possibility of even homodimerization taking place and no possibility of its participation in heterodimerization.

Thirdly, TraesCS3D02G364900 (ABI5) has two attractive acidic-basic electrostatic interactions and four repulsive acidic and basic pairs. This is an indication of heterodimerization of this protein. TraesCS3D02G364900 (ABI5) after one repulsive electrostatic interaction in the second heptad with TraesCS5B02G470600 (EmBP1) is enthralling the repulsive Q↔E’ interaction at *e*↔*g’* positions in the sixth heptad iteratively.

Further, TraesCS7D02G518100 contains an equal number of attractive and repulsive pairs, so it is predicted to form both homo- and heterodimers. We have observed that due to the presence of E↔Q’ at g↔e’ positions in the second heptad position in TraesCS7D02G518100 and TraesCS1A02G072600 pair, there is a possibility of heterodimerization in the sequence. A similar observation in the second heptad is made between TraesCS7D02G518100 and TraesCS7A02G398400, where repulsive E↔E’ pair has been seen at g↔e’ positions with the possibility of strong heterodimerization initiation at second heptad.

TraesCS5D02G124600 (TabZIP1), which has two favorably interacting pairs and five repulsive pairs, is more likely to heterodimerize. Additionally, TraesCS1A02G072600, the most conspicuous member with four repulsive pairs and one attractive pair, is the primary TabZIP with heterodimerization propensity. In the next pair between TraesCS1A02G072600 and TraesCS7A02G398400, repulsive acidic residues are present at g↔e’, indicative of heterodimerization. Furthermore, TraesCS1A02G072600, has repulsive acidic pairs E↔E’ and D↔E’ at g↔e’ positions with TraesCS5B02G470600.

### 3.5 Experimental validation of dimerization properties of bZIP transcription factors in bread wheat

#### A) bZIP TFs protein expression within prokaryotic expression system

Wheat bZIP transcription factors have not been sufficiently characterized at the molecular level due to a lack of cloning and recombinant protein production of wheat bZIPs. Therefore, we started with cloning, and the expression of recombinant wheat bZIP proteins in bacterial system. DNA binding region and leucine Zipper region of bZIPs were amplified as BamHI and HindIII fragments from cDNA templates prepared from respective developmental stages. For the experimental validation of dimerization potential analysis of wheat bZIPs, we cloned seed-specific bZIP TFs in pET23+ vector and expressed in E. coli (BL21 DE3) expression system. All bZIPs have a T7 tag and protein lysate was dialyzed against the low salt buffer. Further, protein lysate was applied to the Heparine-sepharose column and protein was eluted using 1M KCl elution buffer (1M KCl, KPO_4_ buffer, pH 7.4) as shown in Figure 4. Details for further downstream processing and purification steps are provided in the supplementary file S1.

**Figure 4.**
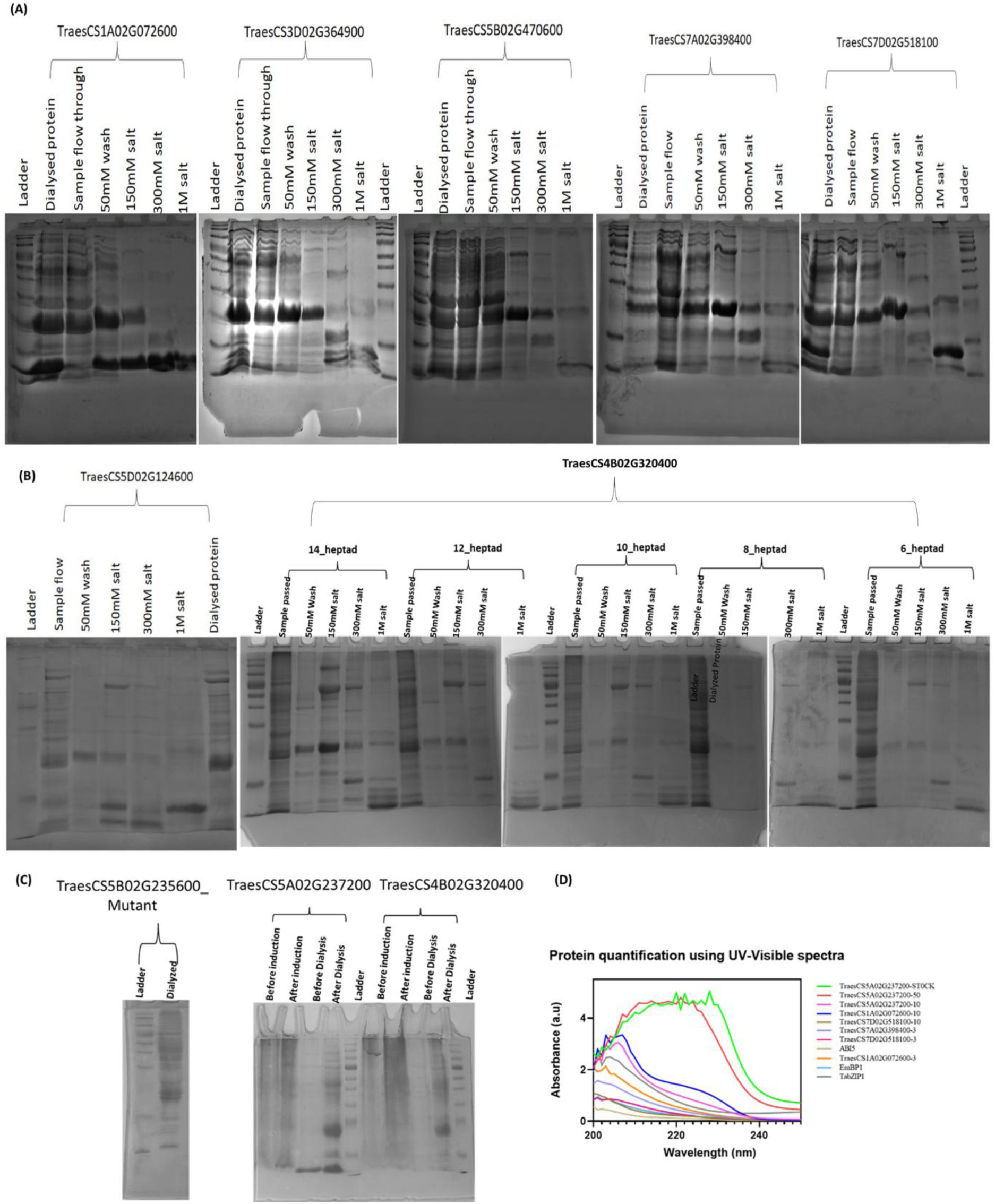
Protein expression of candidate TabZIPs: In panel (A-C) SDS PAGE of expressed TabZIP recombinant proteins in the E-coli bacterial system: TraesCS1A02G072600, TraesCS3D02G364900 (ABI5), TraesCS5B02G470600 (EmBP1), TraesCS7A02G0398400, TraesCS7D02G518100, TraesCS5D02G124600 (TabZIP1), TraesCS5A02G237200, TraesCS5B02G235600 (mutant homeolog), and TraesCS4B02G320400, proteins loaded in the gels following the sequence of expression steps such as Dialysed protein, sample passed through the heparine sepharose column, 50Mm salt wash, 150 mM salt, 300Mm salt and 1M salt elution step for each protein (D) The HPLC-purified proteins were quantified using UV-Visible spectroscopy to measure the concentration for CD and MALDI-TOF MS.

#### 3.5.1 Secondary structure prediction and Thermal stability analysis of TabZIP proteins

To evaluate the thermal stability of all eight candidate TaZIP proteins used in this study, we used circular dichroism spectroscopy at 222 nm to monitor their thermal stability. Figure 5 shows the wavelength scan and temperature-induced melting curves of TraesCS1A02G072600, TraesCS7D02G518100, TraesCS5B02G470600 (EmBP1), TraesCS3D02G364900 (ABI5), TraesCS7A02G398400, TraesCS5D02G124600 (TabZIP1) with 2 µM protein concentration. CD data of TraesCS5A02G237200 (and its mutant TraesCS5A02G23560) and TraesCS4B02G320400 has been shown in Figures 7 and 8 respectively. All proteins showed the cooperative unfolding as the temperature was increased from 6-85°C. Each melting curve was fitted to an equation (equation 1) that describes the equilibrium constant (Kd) as a function of temperature. Assuming the thermal denaturation of leucine zipper domains to be of a two-state type, each melting curve was fitted that gave values of T_m_ and ΔH_m_. Slope of a linear fit of Tm Vs ΔH_m_ gave the value of ΔC_p_ in kcal mol-1K-1(Table 1). This along with Tm and ΔHm values were used to obtain ΔG_D_ at 25°C in Figure 5 A-C panels. All such thermodynamic stability parameters are given in Table 1. Proteins with cysteine residues in the sequence were pre-heated for fifteen minutes with DTT before melting. TraesCS7D02G518100, the most stable transcription factor employed in the study, with the greatest Tm (64°C) and ΔG_D_ value, exhibits exceptional behaviour. Rest of the candidates, such as TraesCS1A02G072600, TraesCS5B02G470600 (EmBP1), TraesCS3D02G364900 (ABI5), TraesCS7A02G398400, TraesCS5D02G124600 (TabZIP1) follow the typical bZIP transcription factor behaviour with Tm range between 36-47°C. Rest two TabZIPs, TraesCS5A02G237200 (and its mutant TraesCS5A02G23560) and TraesCS4B02G320400 considered for the amino acid variations and leucine zipper length features respectively, exhibited Tm ranging from 38°C to a stable Tm 52°C. All thermal denaturation results were observed to be reversible, with Subtle changes in Tm values more/less than 2°C in case of all proteins used in the study except TraesCS5A02G237200 with more changes after heat exposure till 85°C temperature being a homeolog with an unstable thermal behaviour of 30°C.

**Figure 5.**
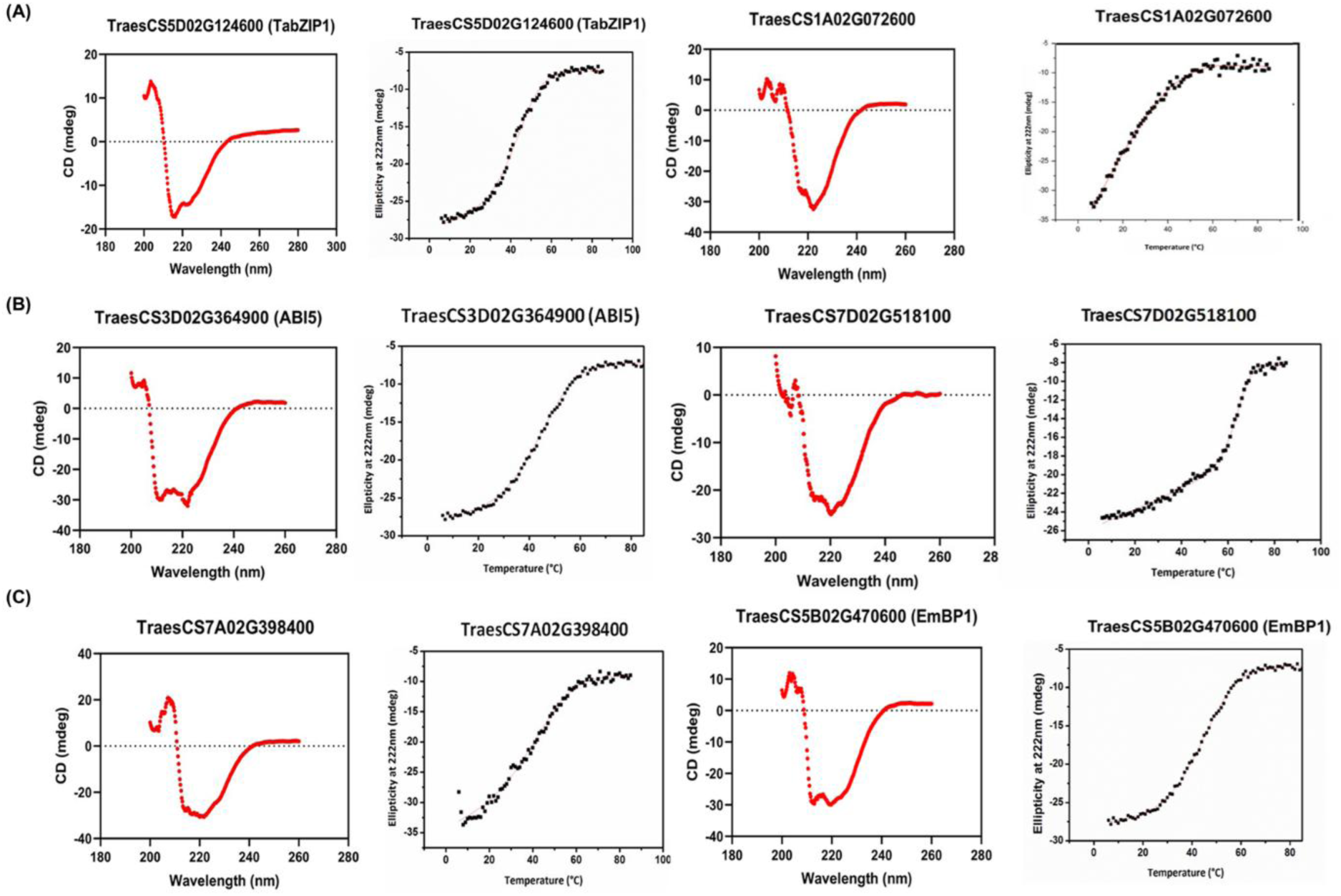
CD wavelength scan and thermal stability analysis of candidate TabZIP proteins. Wavelength scan of bZIP proteins show minima at 222nm and 208 nm, an indicator of alpha helices (red panels). CD thermal denaturation profiles of TabZIPs in 2µM concentration (black panels). The helical content decreased as the temperature was raised from 6-85°C.

**Figure 6.**
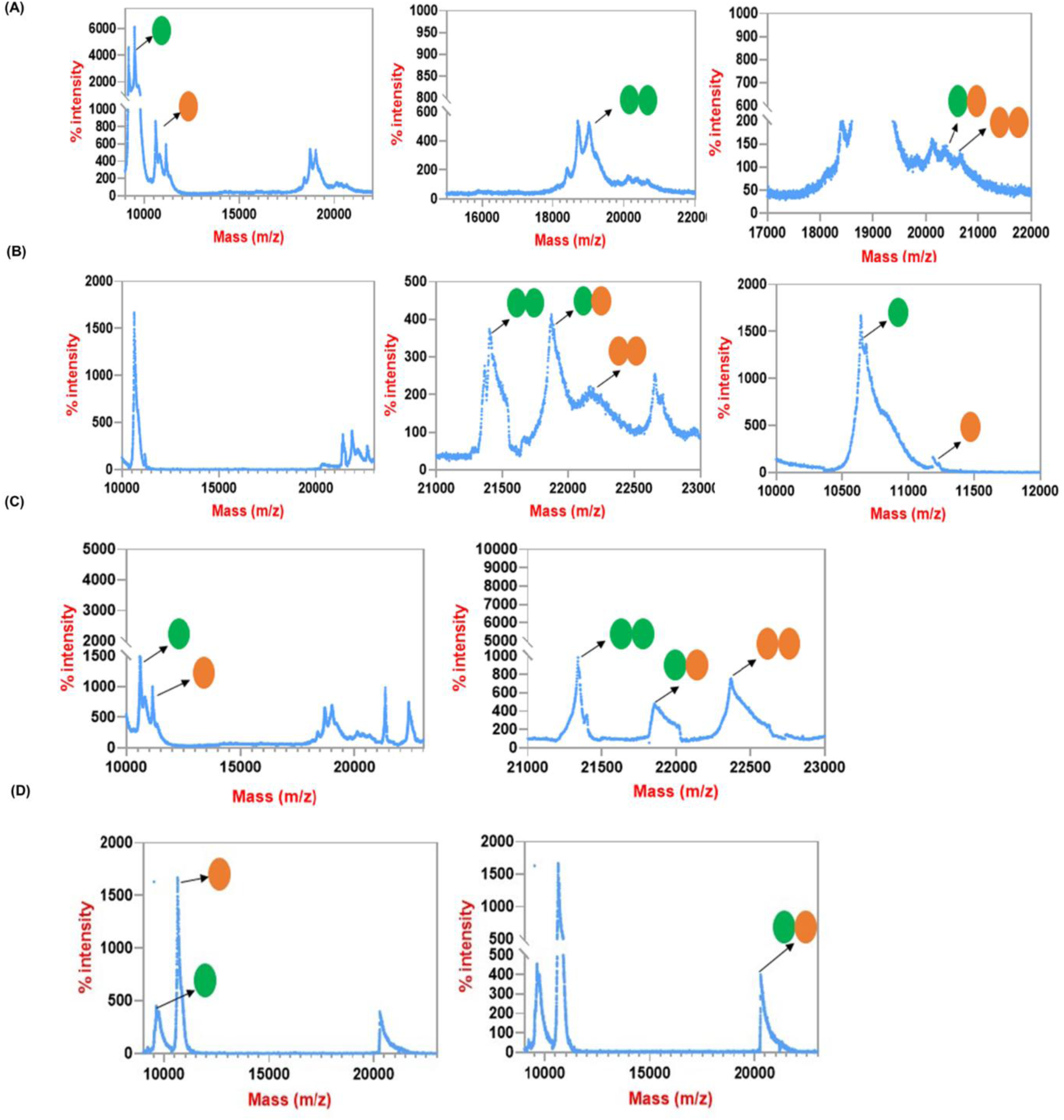
MALDI-TOF mass spectra of candidate TabZIP proteins. (A)TraesCS7A02G398400 and TraesCS3D02G364900 (ABI5) in combination with TraesCS1A02G072600, showing both the monomeric peaks (For better understanding peaks have been labelled with circle shapes; single green or orange coloured) and homo- (two shapes with same colour) and heterodimer (two shapes together with different colours) peaks. (B) MALDI-TOF mass spectra TraesCS1A02G072600 and TraesCS5B02G470600 (EmBP1) showing both the monomeric and dimeric peaks. (C) MALDI-TOF mass spectra of TraesCS5B02G470600 (EmBP1) and TraesCS3D02G364900 (ABI5) showing both the monomeric peaks, homo- and heterodimer peaks. (D) MALDI-TOF mass spectra TraesCS1A02G072600 and TraesCS7A02G398400 showing the monomeric peaks and both the homo- and heterodimeric peaks.

**Figure 7.**
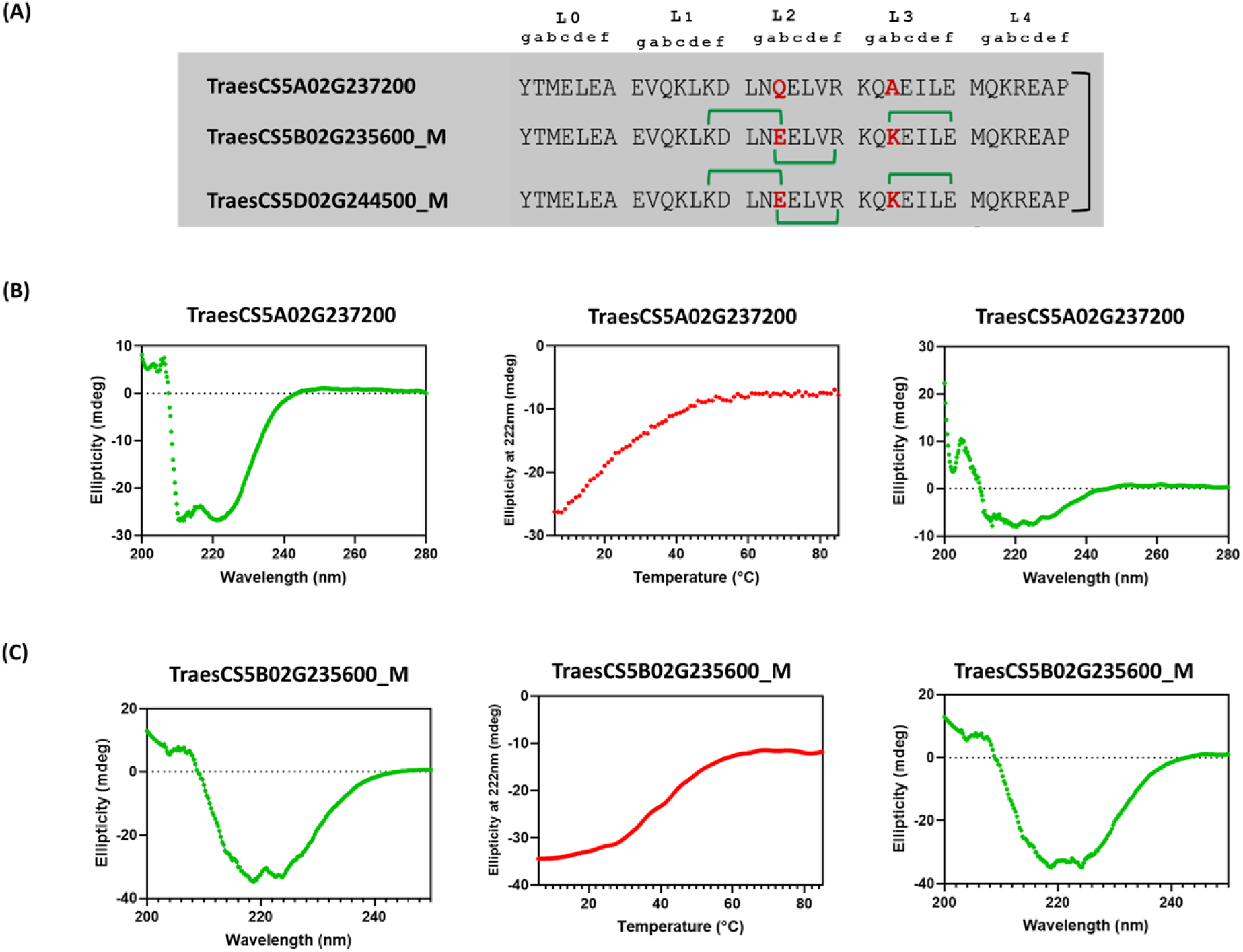
CD wavelength scan and thermal stability analysis of TabZIP TraesCS5A02G237200 and its mutants. (A) Protein sequence alignment of the TraesCS5A02G237200 with its homeologs TraesCS5B02G235600 and TraesCS5D02G244500. (B) and (C) Wavelength scan of bZIP proteins show minima at 222nm and 208 nm, an indicator of alpha helices (left green panels). CD thermal denaturation profiles of TabZIPs in 2µM concentration (red panels). The helical content decreased as the temperature was raised from 6-85°C. Reversibility check was performed by re-scanning the wavelength immediately after the thermal denaturation scan, once the sample had cooled to 6°C (right green panels)

**Figure 8.**
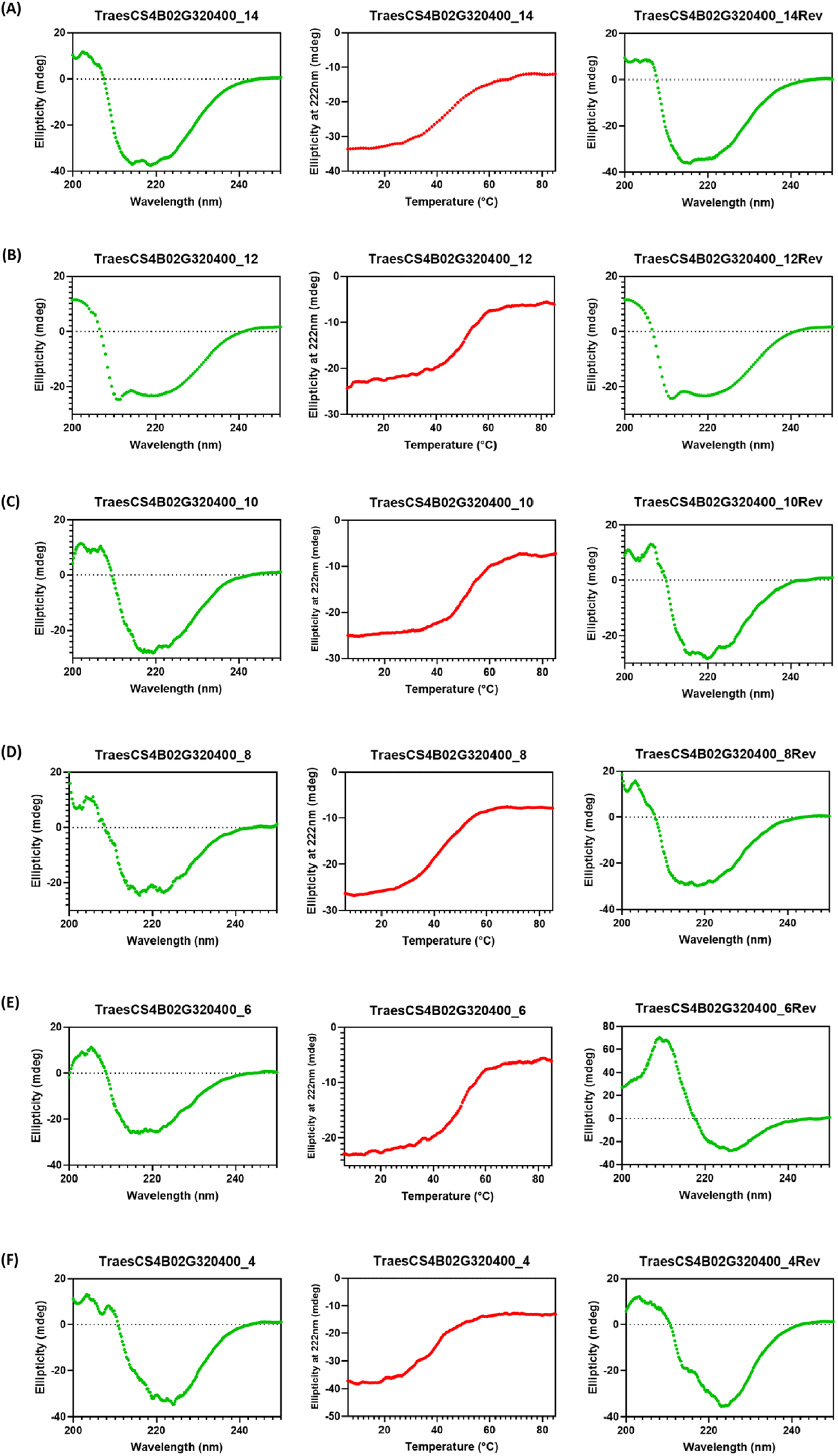
CD wavelength scan and thermal stability analysis of TabZIP TraesCS4B02G320400 and its length variants. (A-F) Wavelength scan of TraesCS4B02G320400 with all length variants, ranging from 4 to 14 heptad repeats, denoted as _4, _6, _8, _10, _12, and _14 heptad lengths, showing minima at 222nm and 208 nm, an indicator of alpha helices (left green panels). CD thermal denaturation profiles of TabZIPs in 2µM concentration (red panels). The helical content decreased as the temperature was raised from 6-85°C. Reversibility check was performed by re-scanning the wavelength immediately after the thermal denaturation scan, once the sample had cooled to 6°C (right green panels)

#### 3.5.2 Protein-protein interaction analysis of TabZIPs

We employed MALDI-TOF to investigate the protein-protein interaction fate or dimer forming capability between the six candidate TabZIPs. It is assumed that a heterodimer between two proteins will be formed only if it is more stable than either of the homodimers in a mixture.

TraesCS7D02G518100 is more stable and shows higher Tm (63°C) and ΔG_D_ values compared to other TabZIPs used in this study. MALDI-TOF MS traces are shown in Figure 6 and the mass of TabZIPs have been provided in Table 1. TraesCS3D02G364900 (ABI5), a well-studied bZIP shows the monomeric monoisotopic peak, and a dimer peak is also observed. TraesCS3D02G364900 (ABI5) contains charged amino acid residues at ‘g’ and ‘e’ positions in the first six heptads. It also may heterodimerize with other bZIP such as TraesCS1A02G072600, TraesCS7A02G398400, and TraesCS5B02G470600 (EmBP1).

Here we show that TraesCS1A02G072600 may heterodimerize with TraesCS5B02G470600 (EmBP1). Supplementary figure 3 shows the MALDI-TOF graph of each mono- and dimeric ion peak for all six TabZIPs considered for the in-vitro protein-protein interaction study. Similarly, TraesCS7A02G398400 dimerizes with TraesCS5B02G470600 (EmBP1). The two TabZIPs such as TraesCS5B02G470600 (EmBP1) and TraesCS3D02G364900 (ABI5), demonstrate dimerization potential not seen in other bZIPs studied here.

### 3.6 Identifying the amino acid substitution and heptad length impact on the dimerization properties in TabZIPs

We tried to extend and validate the earlier gene expression data by structural analysis approach to study the amino acid sequences in TraesCS5A02G237200 (Kaur *et al*. 2024). Different amino acids present in TraesCS5A02G237200 owe additional electrostatic interaction fate to its homeolog (TraesCS5A02G235600) present on the B-subgenome (Figure 7). The higher coupling energy dimension from the literature goes in synergy with our results when we see the occurrence of glutamine (Q) and alanine (A) in TraesCS5A02G237200 with Tm 30°C compared to a significant shift to 43°C in mutant homeologs TraesCS5A02G235600 and TraesCS5A02G244500 having glutamic acid (E) and lysine (K) instead of Q and A in their sequence.

The alignment of 265 protein sequences (Supplementary figure 1) revealed remarkable observations regarding the length of bZIP proteins in wheat compared to human and mouse. Notably, wheat bZIPs tend to be longer, with some extending up to 14 heptads, such as TraesCS4B02G320400. To assess the stability of these extended coiled-coil structures, truncated variants of TraesCS4B02G320400 were designed, ranging from the minimum required 4 heptads (for the helical dimerization between two monomers) to the full 14-heptad length. Thermal denaturation experiments revealed distinct Tₘ values for each variant. The shortest variant (4 heptads) exhibited a Tₘ of 37°C, while longer variants displayed varying levels of stability: 6 heptads (50°C), 8 heptads (44°C), 10 heptads (53°C), 12 heptads (50°C), and 14 heptads (45°C) as shown in Figure 8. The variation in Tₘ across different lengths suggests that the addition of amino acids such as glutamic acid (E), lysine (K), aspartic acid (D), arginine (R) etc. at ‘g’ and ‘e’ positions, influence structural stability, likely due to differences in residue composition and electrostatic interactions. Analysis of the heptad positions in TraesCS4B02G320400 revealed the presence of serine (S), alanine (A), valine (V), isoleucine (I), and cysteine (C) at the ‘a’ position, suggesting a higher potential for heterodimer formation. Notably, a single asparagine (N) was observed in the fifth heptad, which may further indicate homo-dimerization specificity. The diversity of residues at the ‘a’ position indicates possible variations in coiled-coil interactions, contributing to the structural adaptability of the bZIP protein.

## Discussion

Wheat is a low-cost cereal crop. Its grains are a major source of human calories, and wheat straw is used as an animal feed (Awika 2011). Since the draft genome is of Chinese spring cv. *T aestivum*, we proceeded to use the data of this variety. 265 projected bread wheat bZIPs can produce 35, 245 dimers, demonstrating the enormous potential of gene regulation. The DNA binding class of transcription factors follow the dynamic gene expression by occurring at distal ends of chromosomes (Mak *et al*. 2009). In our analysis, a smaller percentage of bZIP transcription factors were detected in the centromeric regions of the chromosomes, whereas the majority were found in the distal ends. This non-random distribution points to a possible cause of expression asymmetry, which is probably impacted by the number of regulatory elements and chromatin accessibility (Smith *et al*. 2011; Perea-Resa and Blower 2018). Furthermore, this pattern might be a reflection of traits linked to the polyploid background, where the genomic organisation of transcription factors is influenced by processes for gene retention and dosage balancing (Conant *et al*. 2014; Cheng *et al*. 2018).

In nature, dimerization is one way the transcription factors increase their repertoire of recognized DNA sequences. Heterodimerization results in the reorganization of novel DNA sequences leading to new functions (Deppmann *et al*. 2004; Rodríguez-Martínez *et al*. 2017). All bZIPs proteins are aligned with the invariable N in the DNA-binding domain of the proteins. Further, the presence of R/K at the eighth position c-terminal to invariable N and L at the thirteenth position, followed by L or any other hydrophobic amino acid at every seventh position or ‘a’ position of a heptad and the presence of charged amino acid at ‘e’ and ‘g’ position regards a sequence as a bona fide bZIP TF. P or two Gs, both natural helix-breakers, establish the bZIP coiled-coil limit at the c-terminal. Out of 294, 29 TabZIPs do not contain a defined bZIP domain and are misclassified under the bZIP family as they contain the conserved bHLH- and DOF-domains in their sequence. We have excluded these transcription factors in our analysis. The alignment of 265 protein sequences led to a few remarkable observations. 1) Compared to human and mouse wheat bZIPs are long. In some cases, they are up to 14 heptads long. Initially, we didn’t anticipate the possibility of such a lengthy coiled-coil to be stable, this hypothesis was also validated in this study. 2) Unlike mammals, wheat bZIPs have a high propensity to form homodimers. 3) Numerous wheat bZIPs feature intricate combinations of amino acids, making it challenging to predict dimerization fate. 4) bZIP homeologs in bread wheat show amino acid substitutions, ranging from single to multiple residue variations due to polyploid nature of evolution. Validation of these sequence differences led to the recognition of homeolog expression switches under different stress and developmental conditions, highlighting the dynamic regulation of homeolog-specific gene expression.

To extend our study on the functional side of TabZIPs, we have expanded the biophysical analysis of these proteins to determine how the particular a, d, e, and g locations relate to dimerization stability. In wheat, we observed that N at ‘a’ position is present in a low proportion (21%) compared to other model plant species i.e., *A thaliana* and other plant species like *Z mays* (26%). We have observed the presence of amino acids S, C, K, A, and Q at ‘a’ position, especially in the initial four heptads, which is uncommon in reference plants such as *Z mays, O sativa, H vulgare and A thaliana*. The amino acids such as S and A are abundant in TabZIPs in instead of N at ‘a’ position. Amino acid S can be phosphorylated; such modification may change the fate of dimerization and DNA binding (Lee *et al*. 2010). Similarly, A is involved in the stress response such as heat and drought environments (Kusano *et al*. 1995).

The α-helical structure is detected in the bZIP proteins by measuring the ellipticity by the CD spectroscopy (Acharya *et al*. 2002). bZIP dimers unfold reversibly and cooperatively to produce monomers in heat and chemical denaturants. The thermal denaturation curves of human bZIPs are well represented by the two-state model allowing the study of individual amino acids contribution to the dimerization specificity (Acharya *et al*. 2002, 2006). In bread wheat, we observed the two-state thermal denaturation curve during the thermal stability analysis for all eight TabZIPs. In addition, we looked for the reversibility trend in the TabZIPs to study the structural changes in the α-helical structure after the temperature exposure from 6 upto 85°C at the rate of 1°C/min by measuring the change in ellipticity at 222 nm wavelength. All eight bZIPs appeared to reverse with quite subtle changes in the ellipticity but TraesCS5A02G237200.

To determine the molecular basis of heterodimer formation between TabZIP proteins, we analyzed their propensities to form dimeric complexes. As described in section 3.1.1, we grouped the bZIPs based on invariable ‘N’, ‘R’ in DNA binding domain and ‘L’ and charged amino acids in the leucine zipper region (Acharya *et al*. 2002). In case of TraesCS5B02G470600 (EmBP1) with TraesCS7D02G518100, TraesCS1A02G072600, and TraesCS3D02G364900 (ABI5) bZIPs, we predicted that based on the presence of N at ‘a’ position in second heptad these proteins should form a homodimer, but the repulsive basic amino acid residues K↔K’, K↔Q’, and K↔K’ at g↔e’ positions in second heptads should encourage heterodimerization. MALDI-TOF studies further experimentally validated these predictions (Figure 6 and Supplementary figure 3).

In Arabidopsis, during seed maturation, it has been shown that the bZIP53 forms a heterodimer complex with the bZIP10, and bZIP25 (Alonso *et al*. 2009; Jain *et al*. 2017). Being polyploid, bread wheat confers the homeolog copies of each gene expressing under the specific developmental and stress response due to genetic and epigenetic reasons. In our earlier research, the genetic basis of the differences in protein sequences of bZIP triads was explored, which may impart structural stability to a specific homeolog enabling the plant to endure the stress conditions better (Kaur *et al*. 2024). Previous research has examined how each amino acid contributes to the specificity of dimerization in humans (Vinson et al. 2002), drosophila (Fassler et al. 2002), and *Arabidopsis* (Deppmann et al. 2004). In addition to these studies, single α-helical domain enriched with amino acid residues such as E, R, K and H, stabilise the α-helices through salt bridges (Steinmetz *et al*. 2007). In line with these findings, we observed a similar trend in TraesCS5A02G237200, which exhibited an unstable thermal denaturation profile with a melting temperature (Tₘ) of 30°C. On the other hand, the mutant homeolog TraesCS5B02G235600 displayed a significantly higher Tₘ of 43°C, indicating increased structural stability. This difference can be attributed to the presence of glutamic acid (E) and lysine (K) replacing the glutamine (Q) and alanine (A) respectively, in the second and third heptad positions of TraesCS5B02G235600, which provide additional electrostatic interactions. These findings emphasize the critical role of charged residues in the dimerization and stable coiled-coil formation, reinforcing the importance of electrostatic interactions in protein stability and function.

Previous studies have identified a maximum of five heptads in human bZIP proteins, suggesting that beyond this length, additional heptads may not significantly enhance dimerization efficiency (Vinson *et al*. 2002). In contrast, our findings in wheat bZIPs reveal extended coiled-coil domains, reaching up to 14 heptads, which raises intriguing questions about the evolutionary and functional significance of these longer sequences. The presence of additional heptads in wheat bZIPs may contribute to increased stability, altered dimerization specificity, or distinct regulatory mechanisms compared to their human counterparts. (Vinson *et al*. 2002). For TraesCS4B02G320400, leucine zipper length was the key criterion to assess its impact on protein behaviour. We hypothesized that beyond 7 to 9 heptads, the helical structure would begin to destabilize. Contrary to our expectations, the variants ranging from 6 to 14 heptads exhibited remarkable stability, with melting temperatures (Tm) ranging from 50–52°C, exceptionally stable compared to the 4-heptad variant, which had a Tm of 38°C. Our findings highlight that the longer leucine zippers, uniquely found in bread wheat compared to other plant species, as well as human and mouse, likely contribute to protein stability. This enhanced stability may be linked to the polyploid nature of bread wheat, providing functional advantages in its regulatory networks (Evans *et al*. 2022).

## Conclusion

In conclusion, creating new wheat varieties with enhanced agronomic features and comprehending the basic biology of polyploid organisms depend on the dimerization potential of each TabZIP protein and asymmetric gene expression in wheat. Contribution of amino acids at specific heptad positions in TabZIPs and the sequence based structural variations in the bZIP homeologs owe specific biochemical features and helps to withstand the stress response by additional electrostatic interaction possibilities within the forming bZIP dimers. Further investigations are necessary to determine whether the extended length of bZIP domains in bread wheat, provides functional advantages in plant-specific transcriptional regulation. Moreover, the insights gained during the study could aid in breeding strategies in crop improvement by manipulating individual or multiple homeologs to modulate trait response.

## Supporting information

Thermodynamic parameters of candidate TabZIPs

## Acknowledgments

RK and VR thank the National Agri-Food Biotechnology Institute (NABI) Executive Director, Mohali, for research facilities and the Department of Biotechnology (DBT), New Delhi, for funding. RK acknowledges the Department of Science and Technology (DST) for the DST-Inspire fellowship.

## Competing Interest

Authors declare no competing interest.

## Conflict of Interest

The authors declare no conflict of interest regarding the research.

## Author contribution

**Raminder Kaur:** Conceptualization, Writing – original draft, Visualization, Validation, Methodology, Investigation, Data curation, Formal analysis, Review & editing. **Nishtha Sharma:** Review & editing. **Prateek Jain:** Review & editing. **Mamta Verma:** Review & editing. **Ashish Apkari:** Review & editing. **Vikas Rishi:** Conceptualization, Methodology, Writing – original draft, Writing – review & editing, Methodology, Investigation, Formal analysis, Supervision, Review & editing, Project administration, Funding acquisition.

## Notes

### Competing Interest Statement

The authors have declared no competing interest.

### Summary of Updates

This version contains an abstract with detailed and improved format which was not clearly described in the earlier submission.

