## Supplementary material for "Mapping the dimerization specificity of bZIP transcription factors in bread wheat": Thermodynamic parameters of candidate TabZIPs

| Protein_id | $T_m$[a], °C | $\Delta H_{m,}$[b], kcal/mol | $\Delta G_D$[c], kcal/mol | Mass (monoisotopic) |
|---|---|---|---|---|
| TraesCS5B02G470600 (EmBP1) | 43 | -38 | -8.9 | 11183 |
| TraesCS5D02G124600 (TabZIP1) | 43.2 | -45 | -9.1 | 12644 |
| TraesCS3D02G364900 (ABI5) | 43.1 | -37 | -8.9 | 10669 |
| TraesCS7A02G398400 | 42.5 | -32 | -8.8 | 10700 |
| TraesCS1A02G072600 | 36.6 | -18 | -8.2 | 9639 |
| TraesCS7D02G518100 | 64 | -87 | -9.5 | 10653 |

**Table 1 Thermodynamic parameters of candidate TabZIPs**

[a]$T_m$ is the midpoint of thermal denaturation. Error in the $T_m$ values of three independent measurements was ≤0.5 °C.

[b] $\Delta H_m$ is the enthalpy change at $T_m$. Values are the mean of three independent measurements and represent ±standard error.

[c]$\Delta G_D$ are the values of the free energy change of dimer formation at 25° C and were calculated using the values of ΔHm at corresponding Tm and a ΔCp value of -1.2 ± 0.13 kcal/ mol/ dimer/ K.
